# Integrin beta 1 and mannose receptor 2 are involved in the antifungal activity of bronchial epithelial cells through *Aspergillus fumigatus* lectin FleA interactions

**DOI:** 10.64898/2026.02.26.708144

**Authors:** N Millet, A. Moreau, M. Tarizzo, L. Marti, A. Varrot, E. Gillon, N. Richard, C. Pionneau, S. Chardonnet, H. Varet, R. Morichon, J. Guitard, L. Guillot, V. Balloy, J. Bigot

## Abstract

*Aspergillus fumigatus* is a world-wide saprophyte filamentous fungus which released conidia, its infectious morphotype, in the atmosphere. These conidia are inhaled daily by humans and can colonize the respiratory tract, where they may develop into hyphae, the invasive morphotype. We previously showed that bronchial epithelial cells (BECs) restrict *A. fumigatus* virulence by inhibiting conidial germination and filament formation through a process requiring PI3K signaling and the conidial fucose-specific lectin FleA. In the present study, we are looking to identify host factors and cellular partners involved in the BEC antifungal response and to define the molecular interactions underpinning FleA recognition.

For this, we analyzed transcriptome of BECs infected with *A. fumigatus* in the presence or absence of the PI3K inhibitor LY294002. Functional involvement of candidate genes was assessed by siRNA knockdown and readouts of fungal filamentation (microscopic scoring and galactomannan release). FleA-interacting host proteins were identified by biotin-FleA affinity co-precipitation coupled to Tandem mass spectrometry, and validated by surface plasmon resonance and biolayer interferometry. The spatiotemporal dynamics of FleA and candidate partners were analyzed by confocal microscopy and proximity ligation assay

We demonstrated that BEC antifungal activity involves at least two complementary pathways: a PI3K/laminin-332 axis promoting conidial adhesion, and a FleA-dependent pathway engaging ITGB1 and MRC2 consistent with lectin uptake and trafficking toward LAMP1-positive compartments. These findings nominate FleA-host receptor interactions as attractive targets for anti-adhesive strategies against *A. fumigatus*.

**Author summary:** Fungal pathogens are an increasing threat to public health, as they are becoming more common and harder to treat due to rising drug resistance. Among them, *Aspergillus fumigatus* has been classified as a critical pathogen by the World Health Organization (WHO). This filamentous fungus delivers spores in the air daily, which are constantly inhaled by humans. In people with weakened immunity, these spores can cause a range of lung diseases known as aspergillosis, with severity ranging from mild to life-threatening. Lung epithelial cells are the first cells of the respiratory tract to encounter inhaled spores. In a previous study, we showed that bronchial cells can prevent spore from developing into filaments, the invasive form of *A. fumigatus* that is responsible for tissue damage. This protective effect depends of on the recognition of a fungal protein called FleA. In the present study, we identified host cell proteins that bind to FleA and transport it into intracellular compartments. Our findings suggest that these proteins help bronchial epithelial cells to internalize fungal spores, thereby blocking their transformation into the invasive filamentous form.

## Introduction

*Aspergillus fumigatus* is an opportunistic fungal pathogen found in the environment. This major fungal pathogen is associated with severe pulmonary infectious and allergic diseases, named aspergillosis that develop in patients with underlying chronic lung diseases or immunosuppressed status (1). Bronchial colonization by *A. fumigatus* can lead to chronic pulmonary aspergillosis in patient with chronic obstructive pulmonary disease or allergic bronchopulmonary aspergillosis in people with cystic fibrosis and participates in the decline of respiratory function. The treatment of aspergillosis has become increasingly challenging due to the limited number of systemically administered antifungal classes, only four, and the rising prevalence of *A. fumigatus* strains resistant to azole derivatives (2–6). The World Health Organization classified *A. fumigatus* in 2022 within the critical group of its Priority Fungal Pathogens List, highlighting the need for prioritizing further research (7).

*A. fumigatus* develops four morphotypes during its growth, ranging from dormant to swelling conidia, followed formation of germinative conidia and then hyphae. Each day, humans inhale 100 to 1000 dormant conidia of *A. fumigatus*, the infectious morphotype released in the air and able to penetrate the respiratory tract due to its small size (2-3 µm) (8). In immunocompetent hosts, innate immune system prevents formation of hyphae, the invasive morphotype. Thus, most inhaled conidia are eliminated by the defence mechanisms of the nasal cavity and upper respiratory tract. Conidia which bypass this barrier and infiltrate the respiratory tract will be first in contact with bronchial epithelial cells (BECs). BECs constitute a physical barrier against inhaled microorganisms and participate in fighting against pathogens through mucociliary clearance and expression of genes related to innate immunity (9).

The interaction of *A. fumigatus* with BECs is related to the capacity of conidia to adhere to cells. *A. fumigatus* as many pathogens, use lectins to mediate pathogen-host interactions (10). Lectins are ubiquitous proteins that discriminate glycan patterns or glycol-epitopes with high selectivity and stereo specificity, allowing them to decipher the information encoded in carbohydrates called glycocode. Lectins-carbohydrates interactions are essential in many biological mechanisms including adhesion, host-cell recognition, signalling pathways, immunity and inflammatory responses (11). In resting conidia, *A. fumigatus* expresses the lectin FleA which is a fucose-specific lectin that mediate adhesion to the epithelial cells via recognition of fucosylated glycoconjugates (12). FleA has been targeted for the development of anti-adhesives glycocompounds as alternative antimicrobial drugs (13–15).

In our previous study, we showed that human BECs were able to express antifungal activity by inhibiting germination of *A. fumigatus* conidia (16). This process was not associated with the release of any soluble components but involved the recognition of the fungal lectin FleA by BECs. Indeed, we demonstrated that exogenous addition of FleA to BECs during *Aspergillus fumigatus* infection abrogated the anti-fungal activity of BECs. PI3 kinase (PI3K) was also involved in this antifungal activity because inhibitors of this kinase decreased significantly the antifungal activity of BECs during *A. fumigatus* infection.

The objective of this study was to better understand the interaction between *Aspergillus fumigatus* spores and bronchial epithelial cells by identifying the cellular receptor of FleA and the protein partners involved in this antifungal process.

## Results

### Genes expression depending of PI3 kinase activation

To identify PI3K pathway-related genes that may contribute to BECs antifungal activity, we performed a transcriptomic profiling of BECs incubated or not with LY294002, a PI3K inhibitor, and infected with *A. fumigatus* for 0, 2 and 4 h. Comparative transcriptomic analysis revealed 3389, 3622 and 4667 genes significantly down-regulated and 2998, 3304 and 4377 genes significantly up-regulated in BECs treated with *vs* without PI3 kinase inhibitor at 0, 2 and 4 h, respectively (**Fig. 1A** and **supplementary Table 1**). We then focused on genes consistently reduced by the PI3K inhibitor across all three time points. Applying additional filters (log2 fold change < 0.66 and expression > 50 reads), we identified 738 commonly down-regulated genes (**Fig. 1B** and **supplementary Table 2**). Among these, we highlighted LAMB3 and LAMC2, which encode the β3 and γ2 chains of laminin-332, a heterotrimeric extracellular matrix glycoprotein composed of α3 (or A3), β3 (or B3) and γ2 (or C2) chains. Laminins are major components secreted of the extracellular matrix and can be exploited by microorganisms, such as fungal conidia, for adhesion thereby promoting colonization and invasion of host tissue (17,18). To validate the impact of PI3K inhibition on laminin B3 and C2 expression, BECs were pretreated with LY294002 1 h prior to with *A. fumigatus* conidia. RT-qPCR analysis showed that *A. fumigatus* infection significantly increased LAMB3 expression after 2 and 4 h, and LAMC2 expression after 4 h. In contrast, in presence of PI3K inhibitor, LAMB3 and LAMC2 were not induced by *A. fumigatus* and their expression remained relatively stable over time (**Fig. 1C** and **1D**). Together, these data indicate that *A. fumigatus-*induced expression of laminin-332 is PI3K-dependent, supporting a role for the PI3K pathway in the epithelial antifungal response.

**Figure 1:**
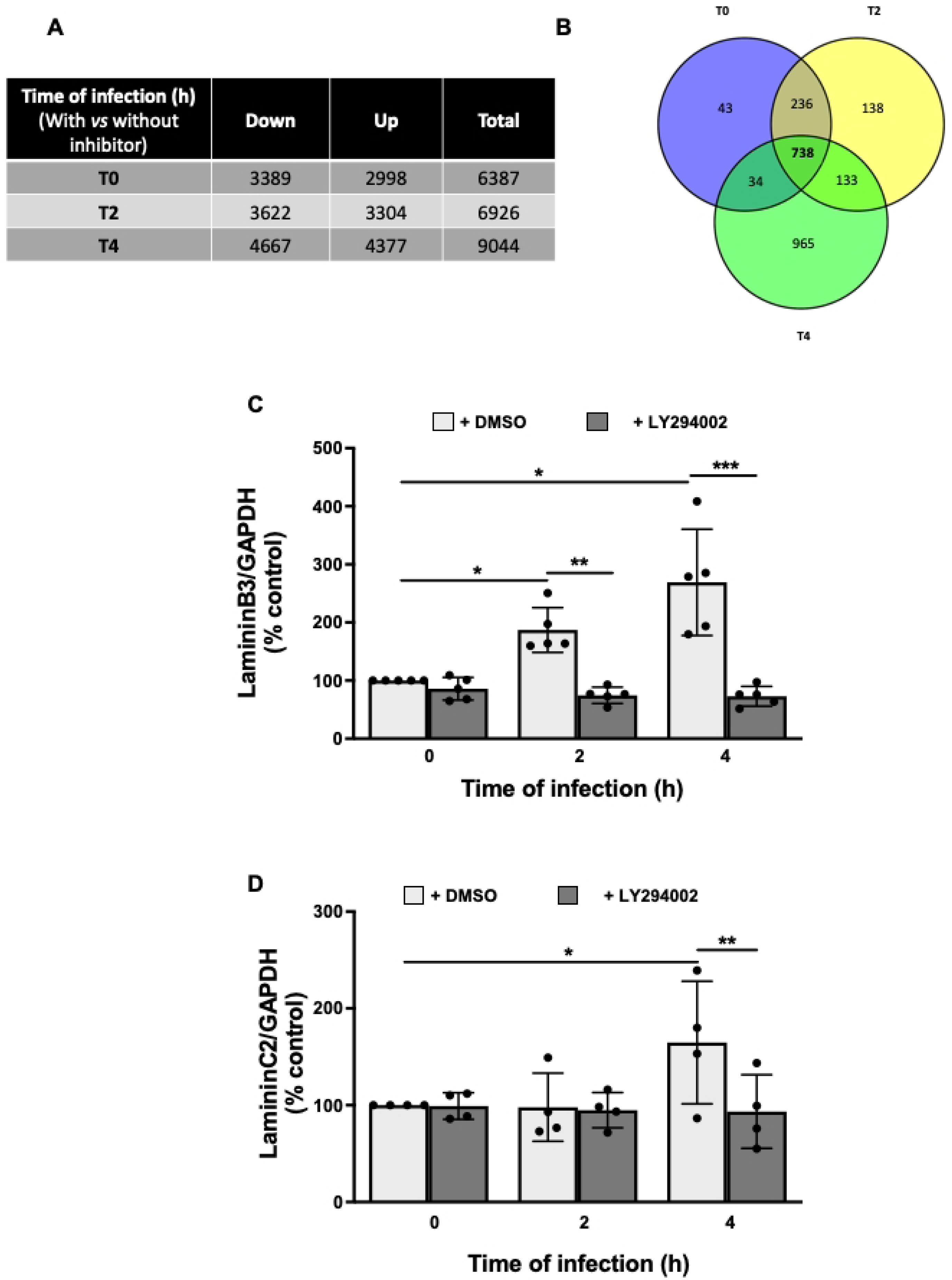
Transcriptomic analysis of BECs preincubated with PI3 kinase inhibitor and then infected by *A. fumigatus*. (**A**) Results of the transcriptomic analysis of human bronchial epithelial cell line with or without inhibitor of PI3 kinase and infected with *A.fumigatus* during 2 h (T2) and 4 h (T4) or not infected (T0). p-value adjustment BH <0.001. (**B**) Classification of the significantly down-regulated genes during the infection time (Log2Foldchange < 0.66 and > 50 reads). Effect of PI3 kinase inhibition on mRNA expression of Laminin B3 (**C**) and laminin C2 (**D**) quantified by qPCR and compared to expression of GAPDH. % of control (T0): epithelial cells non infected nor treated with PI3 kinase inhibitor (LY294002 - 30 µM). Data are presented as mean ± SD, n = 4-5 independent experiments. * p < 0.05, ** p < 0.01, *** p < 0.001 (ANOVA test followed by Bonferroni’s multiple comparison test)

### Involvement of laminin-332 in antifungal activity of BECs

To investigate the involvement of laminin in the antifungal activity, BECs were transfected, for 48 h, with siRNAs targeting LAMB3 and LAMC2. This resulted in a significant knockdown of LAMB3 and LAMC2 expression by 77.1% and 87.1%, respectively, compared with control siRNA-transfected cells (**Fig. 2A**). Confluent transfected BECs were then infected with *A. fumigatus* conidia for 15 h to assess fungal filamentation. Silencing LAMB3 or LAMC2 individually did not measurably affect filament formation (data not shown). In contrast, simultaneous knockdown of LAMB3 and LAMC2 led to a marked increase in filamentation compared with control cells (100 ± 19 *vs* 131.9 ± 13.8, respectively) (**Fig. 2B**). This phenotype was confirmed by quantifying galactomannan released by *A. fumigatus* into the supernatant during filament formation. The galactomannan index increased significantly from 100.0 % ± 11.1 in control cells to 125.2 % ± 24.2 in BECs transfected with LAMB3/LAMC2 siRNA (**Fig. 2C**). Together, these data indicate that reducing LAMB3 and LAMC2 expression impairs the antifungal activity of BECs. To further validate the involvement of laminin-332 in antifungal activity, BECs were pre-incubated for 1 h with increasing concentrations of exogenous laminin 332, and then infected with *A. fumigatus* for 15 h. Galactomannan quantification showed a significant dose-dependent decrease when *A. fumigatus* infected BECs in presence of exogenous laminin 332 (**Fig. 2D**).

**Figure 2:**
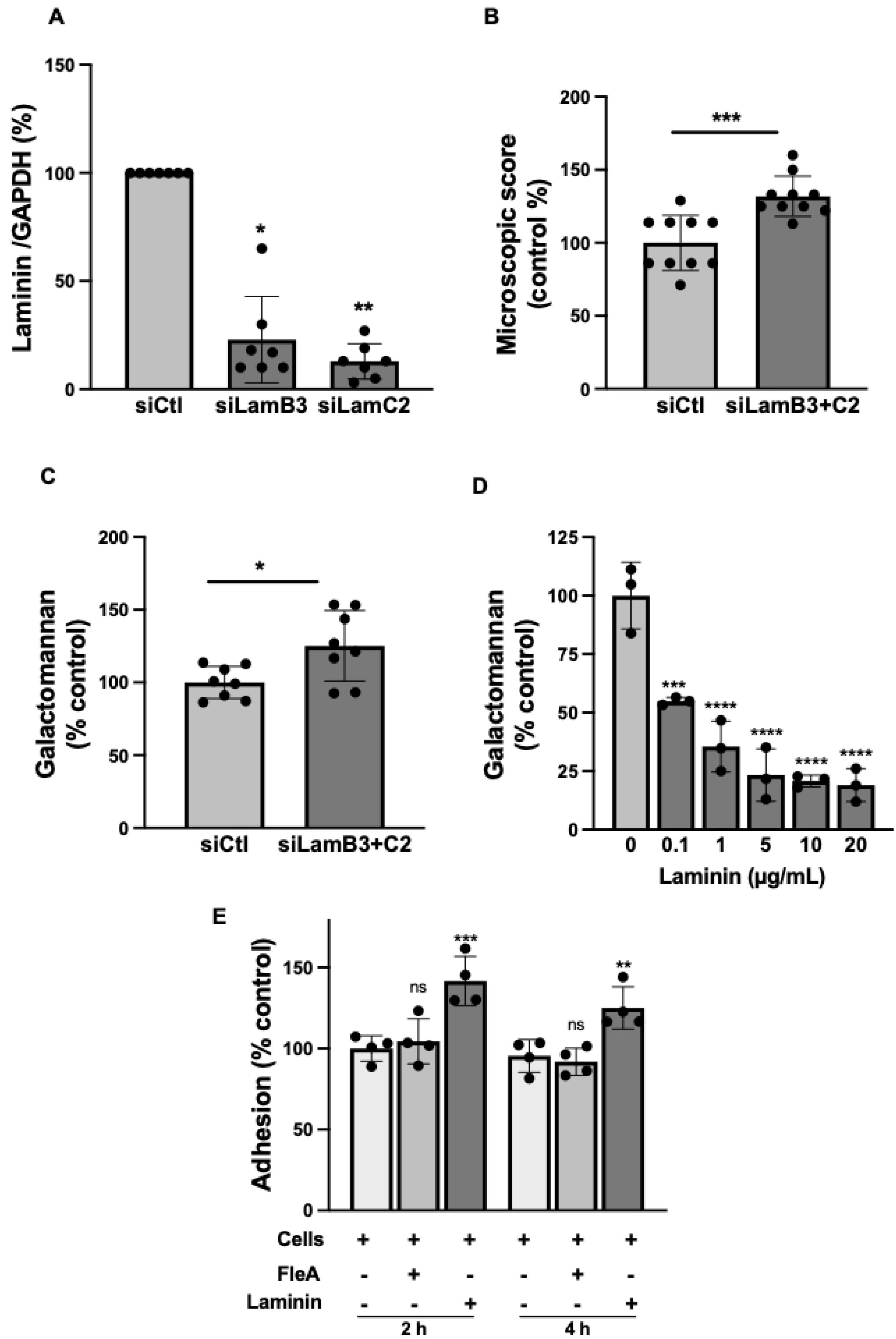
Validation of laminin-332 implication in the antifungal process. (**A**) mRNA Laminin B3 and C2 expression after BECs transfection with siRNA-LAMB3 or siRNA-LAMC2. Data are expressed as percentage of control (siCtl) and presented as mean ± SD, n = 7 independent experiments. * p < 0.05, ** p < 0.01 (ANOVA test). (**B)** Microscopic score of *A. fumigatus* filament formation after infection of BECs transfected with siRNA-LAMB3 and siRNA-LAMC2. BECs were transfected 48 h then infected for 15 h. Data are expressed as percentage of control (siCtl) and presented as mean ± SD, n = 10 independent experiments. *** p < 0.001 (ANOVA test). (**C**) Measure of galactomannan released in supernatant of infected BECs transfected with siRNA-LAMB3 or siRNA-LAMC2. Data are expressed as percentage of control (siCtl) and presented as mean ± SD, n= 8 independent experiments. * p < 0.05 (Student’s t-test). (**D**) Measure of galactomannan released in supernatant of BECs infected with *A. fumigatus* in presence of different concentrations of exogenous laminin-332. Data are expressed as percentage of control (0 µg/mL laminin-332) and presented as mean ± SEM, n = 3 independent experiments. * p < 0.05 (ANOVA test followed by Bonferroni’s multiple comparison test). **(E)** Conidia adhesion on BECs cells after 1 h of pre-incubation with laminin-332 (5 µg/mL) or FleA lectin (2 µM) and infection with 300 conidia/well during 2 h or 4 h. Control: Conidia adhesion to non-treated BECs. Data are expressed as % of control and presented as mean ± SD, n = 4 independent experiments. ** p < 0.01, *** p < 0.001 (ANOVA test followed by Bonferroni’s multiple comparison test).

Thus, supplementation with laminin-332 enhances the antifungal activity of BECs, supporting a functional role of laminin-332 in this process.

### Role of laminin-332 and FleA in conidial adhesion

*A. fumigatus* conidia are known to interact with host extracellular matrix proteins such as laminin (19). To distinguish the role of laminin 332 from that of FleA in the BEC antifungal activity, we investigated their involvement in conidial adhesion. BECs were preincubated for 1 hour with either exogenous laminin 332 or FleA and the treatment was maintained during infection with *A. fumigatus* conidia for 2 or 4 h. Exogenous laminin-332 significantly increased conidial adhesion, reaching 142% ± 15% at 2 h and 125% ± 13% at 4h compared with control cells (100% ± 8%). In contrast, FleA incubation did not alter conidial adhesion to BECs (**Fig. 2E**).

### Identification of cellular protein partners involved in FleA lectin recognition

We next sought to identify the BEC receptors that bind the *A. fumigatus* lectin FleA, by performing affinity co-precipitation by using biotinylated FleA lectin. To ensure that the biotin tag addition to FleA did not affect its binding to BECs and the resulting antifungal activity, cells were pre-incubated for 1 h with either biotinylated or not unmodified FleA, and during the 15- h infection with *A. fumigatus*. As previously reported, we observed a restored filament formation whether FleA was tagged or not, this observation was further supported by galactomannan quantification which showed no significant difference in galactomannan index whether FleA was biotinylated or unmodified (**Supplementary Fig. 1**). To perform co-precipitation, BECs were incubated with biotinylated FleA for 2 h, lysed, and FleA-associated proteins were analyses by LC-MS/MS (**Supplementary Table 3**). The same procedure was performed with non-tagged FleA served as negative control, to exclude non-specific binding. We then focused on the ten most abundant candidate proteins: integrin β1 (ITGB1), catenin beta-1 (CTNB1), mitochondrial trifunctionnal protein α subunit (ECHA), C-type mannose receptor 2 (MRC2), prohibitin-2 (PHB2), synaptic vesicle membrane protein VAT-1 homolog (VAT1), lysosome-associated membrane glycoprotein-1 (LAMP1), prolow-density lipoprotein receptor-related protein 1 (LRP1) and prohibitin (PHB) (**Fig. 3A**). KEGG pathways enrichment analysis indicated that three of the coprecipitated proteins, ITGB1, MRC2 and LAMP1, mapped to the phagosome, the only significantly enriched pathway (**Fig. 3B**). Finally, interaction network analysis using the STRING database (https://string-db.org/) identified predicted associations among these phagosome-related candidates, highlighting a set of three proteins likely to interact with one another (**Fig. 3C**).

**Figure 3:**
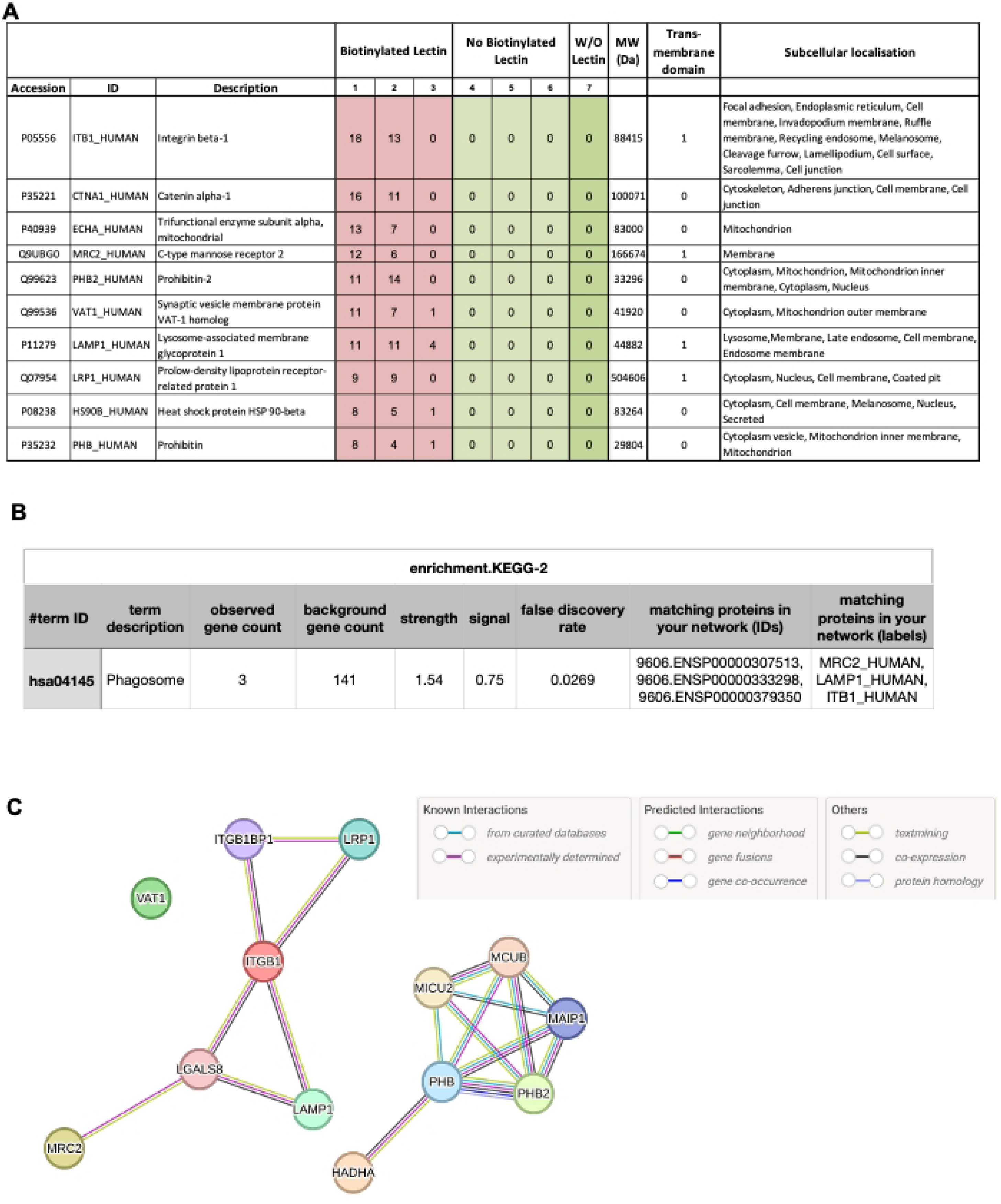
Classification of FleA coprecipitated proteins. Ten most abundant proteins coprecipitated with FleA biotinylated **(A)**. Classification through KEGG pathways enrichment **(B)**. Known and predicted protein-protein interactions using the STRING database. Colored nodes represent query proteins and first shell of interactors **(C)**.

### Role of ITGB1, MRC2 and LAMP1 in the BECs antifungal process

To assess whether ITGB1, MRC2 and LAMP1 contribute to the antifungal activity of BECs, we silenced each gene by siRNA transfection, 72 h prior to *A. fumigatus* infection. Knockdown efficiency was evaluated by RT-qPCR showing that mRNA expression of ITGB1 was inhibited by 93.7% ± 2.2% and 87.0% ± 10.4% when BECs were transfected with siITGB1 or siITGB1+MRC2, respectively. mRNA expression of MRC2 was inhibited by 58% ± 12,2% and 73.6% ± 8% when BECs were transfected with siMRC2 or siITGB1+MRC2, respectively. LAMP1 expression was inhibited by 91.8 ± 7.5% when BECs were transfected with siLAMP1 (**Supplementary Fig. 2**). Microscopic examination indicated that silencing ITGB1 or MRC2 restored fungal filament formation, whereas LAMP1 silencing had no detectable effect. These observations were corroborated by galactomannan measurement in culture cells supernatants which was increased from 100 % ± 21 % in infected BECs transfected with siControl to 171.8 % ± 39.4 % and 230.5 % ± 50.7 %, when *A. fumigatus* infected cells were transfected with siRNA of ITGB1 and MRC2, respectively. In contrast, LAMP1 silencing did not significantly alter galactomannan release (109.0% ± 27.6%) compared to siControl. Notably, combined inhibition of ITGB1 and MRC2 did not further increase filamentation beyond MRC2 knockdown alone (**Fig. 4**).

**Figure 4:**
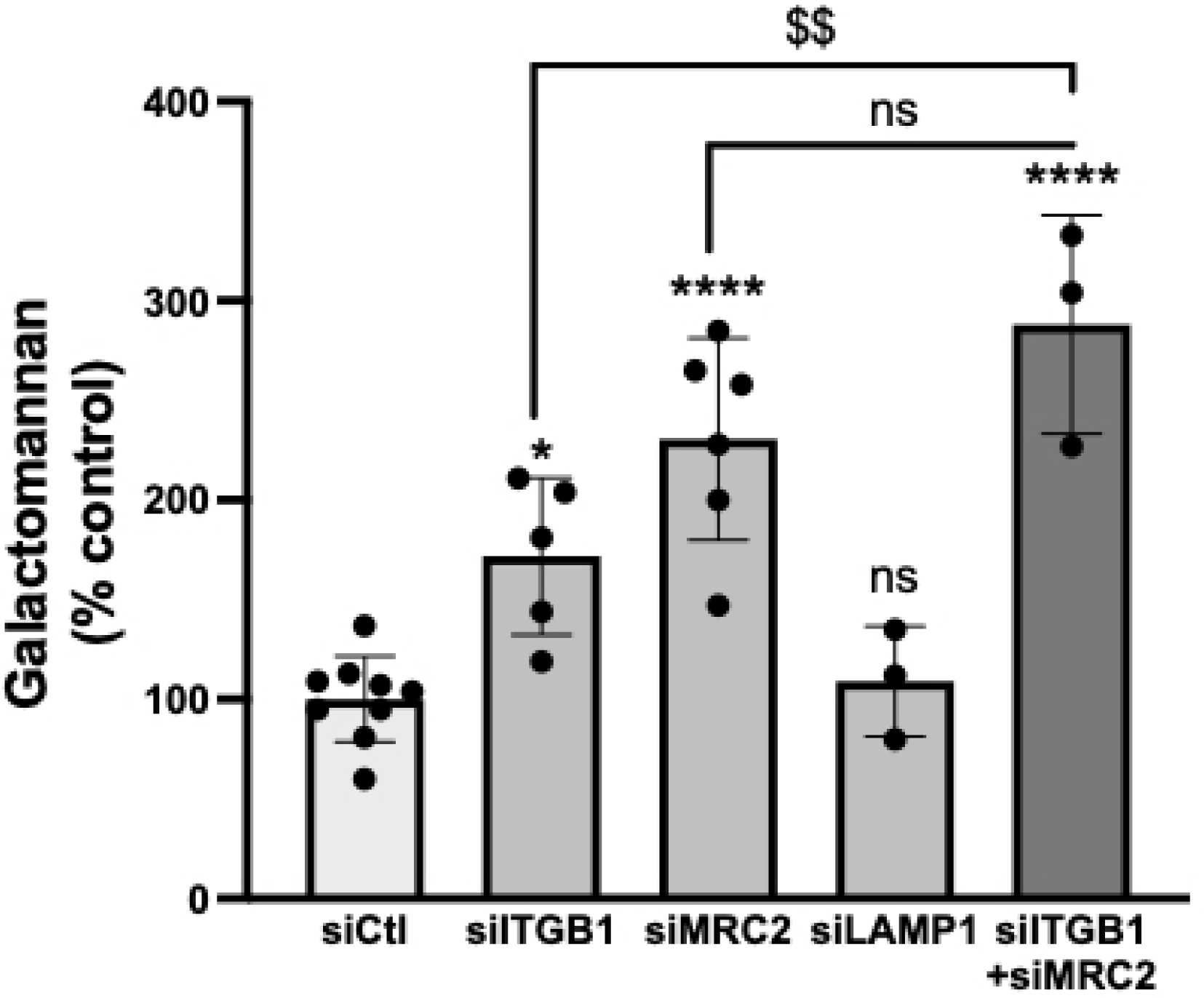
Role of ITGB1, MRC2 and LAMP1 in antifungal activity. Measurement of galactomannan released in BECs transfected with specific ITGB1, MRC2, LAMP1, ITGB1+MRC2 or control siRNA for 72 h and then infected with *A. fumigatus* conidia for 15 h. Data are expressed as % of control and presented as mean ± SD, n = 3-6 independent experiments. *p < 0.05, **** p < 0.0001 (ANOVA test followed by Bonferroni’s multiple comparison test)

Finally, we examined our transcriptomic dataset to determine whether PI3K inhibition affected ITGB1, MRC2 and LAMP1 expression. None of these genes was significantly modulated by *A. fumigatus* infection, nor were they significantly down-regulated by PI3K inhibitor (data not shown).

### Analysis of the interaction of ITGB1 and MRC2 with FleA

To confirm FleA interaction with ITGB1 and MRC2, the two major coprecipitated proteins expressed at the cellular membrane and involved in the antifungal process, we measured binding with FleA using surface plasmon resonance (SPR) and biolayer interferometry (BLI). For MRC2, we used a truncated recombinant construct (aa 31-530) that contains two of the seven C-type lectin domains and three out of its seven glycosylation sites accounting 20% of the protein’s molecular weight. ITGB1 is also highly glycosylated (13 sites) with sugars contributing to almost one-third of its molecular weight. When FleA was immobilized on the biosensors, we obtained for ITGB1 an apparent K_D_ of 5.6 +/- 1.8 nM by SPR and 33 +/- 3.6 nM by BLI after steady state analysis (**Fig 5A** and **B**). We also repeated the experiment in BLI where IGTB1 and MRC2 where immobilised on a Ni-NTA sensor thanks to their His-tag. As again the sensor could not be regenerated fully either with fucose or Glycine pH 1.7 as recommended, single cycle kinetic serie measurements were done giving an apparent k_D_ of 9.9 +/- 2.7 nM and 4.8 +/- 1.4 nM for IGTB1 and MRC2, respectively (**Fig 5C** and **D**). Despite some variations in the values obtained by the different setups and technic, the apparent dissociation constant obtained are coherent and support nanomolar-affinity binding of FleA to both ITGB1 and MRC2.

**Figure 5:**
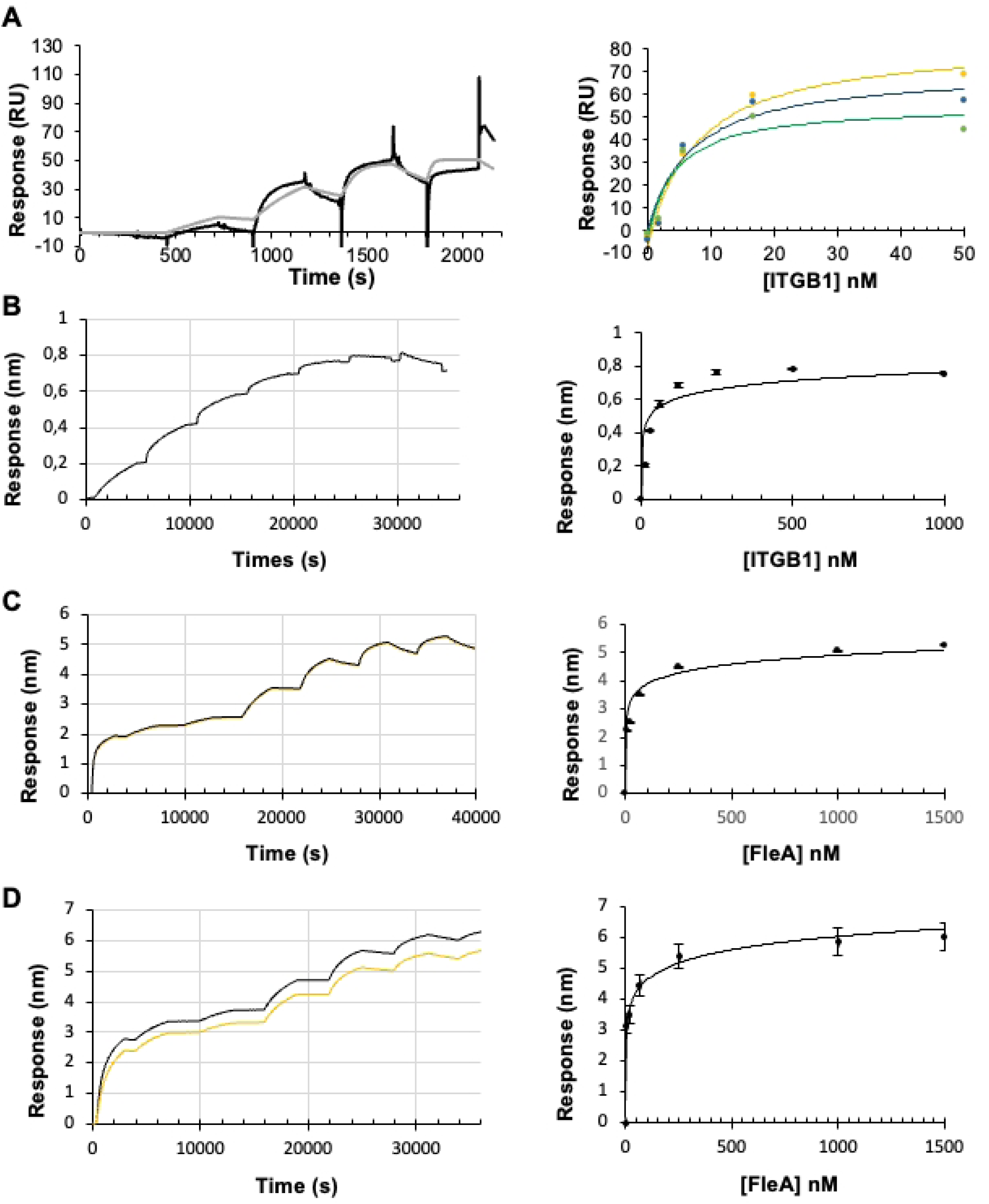
Determination of FleA recognition of ITGB1 or MRC2 by SPR and BLI. **(A**) Representative sensorgram representing binding of ITGB1 (analyte) to immobilized biotinylated FleA (ligand) in SPR kinetic titration serie for concentrations of 1.85, 5.5, 16.6 and 50 nM (left) and the steady-state affinity fit (right) where raw data are represented as points with the corresponding fitted curve colored for each experiment. (**B**) Representative sensorgram representing binding of ITGB1 (analyte) to immobilized biotinylated FleA (ligand) in BLI kinetic titration serie for concentrations of 15.6, 31.3, 62.5, 125, 250, 500 and 1000 nM (left) and the mean steady-state affinity fit with standard deviation on the duplicate (right). Sensorgrams for the duplicate BLI kinetic titration after subtraction of the reference representing binding of FleA (analyte) to immobilized His tagged ITGB1(**C**) or MRC2 (**D**) for concentrations of 3.9, 15.65, 65.5, 250, 1000 and 1500 nM (left) and mean the steady-state affinity fit with standard deviation of the duplicate (right).

### Kinetics of FleA interaction with ITGB1, MRC2 and LAMP1

We next examined, *in situ* by confocal microscopy, the temporal dynamics of FleA associations with ITGB1, MRC2 by incubating BECs with FITC-labelled FleA for 15 and 30 min and for 1, 2 and 4 h. Although LAMP1 silencing did not alter antifungal activity, LAMP1 is a lysosomal membrane protein associated with internalized *A. fumigatus* conidia (20), prompting us to investigate whether it contributes to intracellular trafficking of the lectin. At baseline (non-stimulated conditions), ITGB1, MRC2 and LAMP1 were detected both at the plasma membrane and in the cytoplasm (**Fig. 6A**, **B** and **C**). Upon FleA addition, we observed rapid lectin uptake, as early as 15 min (**Fig. 6A**, **B** and **C**). ITGB1 showed transient co-localization with FleA at 15 and 30 min, initially at the membrane and more prominently in the cytoplasm (white signal) but this association was no longer evident at later time points (**Fig. 6A**). In contrast, FleA exposure markedly increased MRC2 localization at the plasma membrane at15 and 30 min (**Fig. 6B**). Strong cytoplasmic co-localization between MRC2 and FleA was detectable from 15 min and persisted through 4 h (white signal). Notably, after 2 h incubation with FleA, MRC2 displays a distinct redistribution, forming intracellular aggregates that co-localized with the lectin (white puncta, **Fig. 6B**). Consistent with this observation, aggregates quantification showed a significant increase at 2 and 4 h in the presence of FleA (**Fig. 6D**). LAMP1 exhibited robust cytoplasmic co-localization with FleA, at all-time points, and at 2 and 4 h, formed punctate structures with FleA consistant with trafficking toward lysosome compartments (**Fig. 6C**). the number of FleA-LAMP1 aggregates was significantly increased at the 4 h compared to baseline (**Fig. 6E**). Overall, these data indicated that ITGB1 associates with FleA at early time points (15-30 min), whereas MRC2 and LAMP1 display sustained co-localization with FleA throughout the time course, supporting a role in lectin uptake and intracellular trafficking.

**Figure 6:**
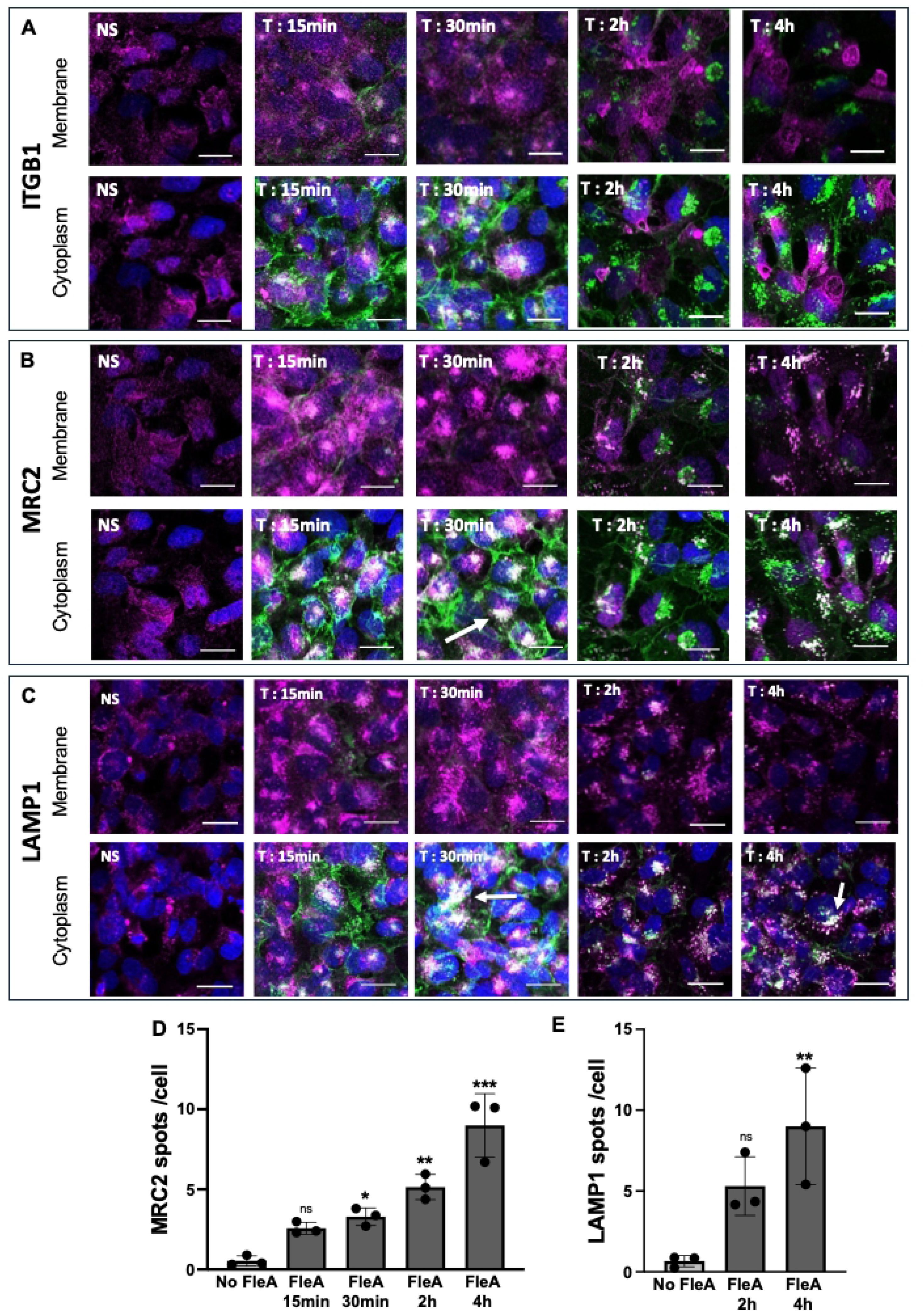
Confocal microscopic analysis of FleA, ITGB1, MRC2 and LAMP1 localization. BECs stimulated with FITC FleA (green) for 15 min, 30 min, 2 h and 4 h or non-stimulated (NS), fixed and stained for ITGB1 (pink) **(A)**, MRC2 (pink) **(B)** and LAMP1 (pink) **(C).** The merge fluorescence between FleA and ITGB1 or MRC2 or LAMP1 is visualized in white. Quantification of white spots (white arrows) visualizing overlay fluorescence (merge) of MRC2 and FleA **(D)** and LAMP1 and FleA **(E)**. Data are expressed as number of white puncta per cell and presented as mean ± SD, n=3 independent experiments. *p < 0.05, ** p < 0.01, *** p < 0.001 (ANOVA test followed by Bonferroni’s multiple comparison test)

### Interactions between ITGB1, MRC2 and LAMP1 in BECs incubated by FleA

To investigate potential associations among ITGB1, MRC2 and LAMP1, we performed an *in situ* proximity ligation assay (PLA) in BECs incubated with FITC FleA for 15 and 30 min, and 1 and 4 h. In both unstimulated cells and FleA-treated cells, we did not detect PLA signals for ITGB1-MRC2 (**Fig. 7A**) or ITGB1-LAMP1 (**Fig. 7B**), indicating no measurable proximity between these pairs under our conditions.

**Figure 7:**
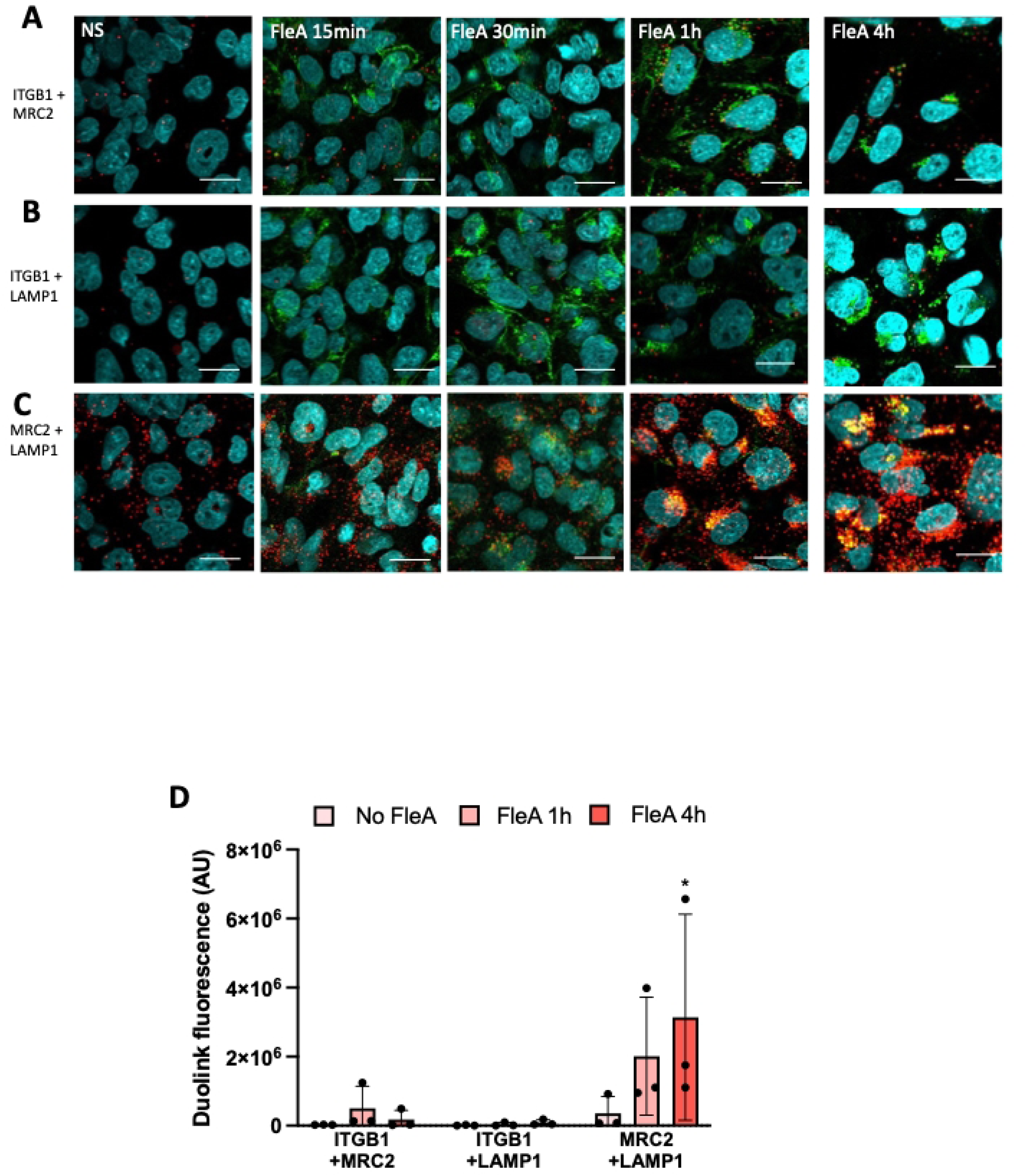
Detection of ITGB1-MRC2, ITGB1-LAMP1 and MRC2-LAMP1 interactions in BECs stimulated by FleA. Proximity ligation assay permitting detection of protein-protein interactions in situ (<40 nm distance) at endogenous protein levels was performed in BECs stimulated by FleA for 15 min, 30 min 1 hour and 4 h or not stimulated (NS). Specific ITGB1, MRC2 and LAMP1 antibodies were then linked covalently to specific DNA primers. A hybridization step followed by a PCR amplification with fluorescent probes then permits visualization of ITGB1-MRC2 **(A)**, ITGB1-LAMP1 **(B)**, MRC2-LAMP1 **(C)** interactions as red spots of proximity by fluorescence microscopy. Quantification of red spots visualizing protein-protein interactions is expressed as arbitrary units and presented as mean ± SD, n=3 independent experiments. *p < 0.05 (ANOVA test followed by Bonferroni’s multiple comparison test) **(D)**.

In contrast, MRC2-LAMP1 interactions were barely detectable at baseline but increased markedly upon FleA stimulation, becoming evident after 1 h and reaching significance at 4 h (red PLA puncta; **Fig. 7C**). Moreover, the MRC2-LAMP1 PLA signal showed strong co-localization with FITC-FleA at 1 and 4 h (yellow signal), supporting the formation of a FleA-associated MRC2-LAMP1 complex. This was confirmed by fluorescence quantification (**Fig. 7D**).

## Discussion

In our previous study, we showed that BECs can attenuate *A. fumigatus* virulence by inhibiting conidial germination, limiting filaments formation and thereby restricting tissue invasion. We also demonstrated that this epithelial antifungal response involves the PI3K pathway and the fungal lectin FleA (16). In the present study, we identify, for the first time, three host factors implicated in this process, laminin-332, ITGB1 and MRC2, and describe their temporal involvement.

The first step for infection or invasion is the attachment of microorganisms to host extracellular (ECM) components such as laminin and collagen. Several *A. fumigatus* proteins have been reported to bind laminin, although the specific laminin isoform was not defined. Thus, Gil *et al*. described a 37-kDa conidial polypeptide that binds cellular laminin (19), and Tronchin *et al*. reported that laminin binding at the conidial surface is swelling-dependent and mediated by a 72-kDa-cell-wall glycoprotein (21). Laminin also interacts with the recombinant protein, Aspf2, a major allergen expressed by *A. fumigatus* mycelium (22), through the extracellular thaumatin-domain protein AfCalAp, and antibody-based experiments further suggest its surface expression on swollen conidia (23).

Laminins are heterodimeric ECM glycoproteins composed of one α, one β and one γ chain and are named according to their subunit compositions. The three chains are expressed independently and assembled into trimers prior to secretion. In our model, *A. fumigatus* infection of BECs increased expression of LAMB3 and LAMC2, two chains that together define laminin-332 which is constituted of α3, β3 and γ2 chains. Notably, the infection-induced upregulation of LAMB3 and LAMC2 was abolished by PI3K inhibitor, LY294002, indicating that laminin-332 induction is PI3K-dependent in this context. The involvement of PI3 kinase in the laminin 332 expression has been reported in other cellular models. Transcriptomic analyses in oesophageal epithelial cells indicated that laminin-332 synthesis depends on PI3K activity and that laminin can, in turn, activate PI3K signaling (24). Similarly, IL-1β-induced laminin-332 expression in islets of Langerhans was inhibited by LY294002 (25). Here, we demonstrated that laminin-332 contibutes to epithelial antifungal process. Indeed, simultaneous silencing of LAMB3 and LAMC2 in BECs increased *A. fumigatus* filament formation and elevated galactomannan release in infected-cells supernatant, consistent with a reduction of BECs antifungal activity. The role of laminin 332 in the antifungal process was reinforced by the addition of exogenous laminin 332 to the *A. fumigatus* infected BECs that increased inhibition of the filament formation. Moreover, addition of exogenous laminin 332 enabled to determine its role in the conidial adhesion to BECs by showing a significantly increase of conidial adhesion. Altogether, these data indicate that *A. fumigatus* infection induces laminin-332 expression by BECs in a PI3K-dependent manner and that laminin-332 promotes conidial adhesion to the epithelial surface.

In parallel, affinity co-precipitation identified three FleA-associated host proteins, ITGB1, MRC2 and LAMP1, all mapping to the phagosome pathway. Notably, laminin-332 was not detected among the FleA co-precipitated proteins, and transcriptomic analyses indicated that ITGB1, MRC2 and LAMP1 expression is not PI3K -dependant under our conditions. Functional assays further showed that ITGB1 and MRC2 contribute to antifungal activity, as their silencing increased fungal filamention, whereas LAMP1 knockdown did not affect filament inhibition, suggesting that LAMP1 is not essential for this readout. Together, these findings support the existence of at least two separable components with the epithelial antifungal response: a PI3K-dependent laminin-332 axis and a FleA-associated ITGB1/MRC2 axis.

Direct binding of FLeA to ITGB1 and MRC2 was confirmed by SPR and BLI, revealing nanomolar apparent affinities. Given that the affinity of FleA for fucose of fucosylated oligosaccharides ranges between 50 and 150 µM, these results provide a clear illustration of how multivalency increases avidity in lectin–glycan interactions. The ITGB1 and MRC2 proteins carry multiple N-glycosylation sites, and dimeric FleA is dodecavalent with each protomer presenting 6 binding sites but recognition of a single glycosylation site would be sufficient for the interaction (12, 26). Consistent with sugar-dependent recognition, partial regeneration of biosensors could be achieved with fucose, whether FleA is used as ligand or analyte.

Finally, confocal microscopy revealed distinct kinetics of FleA associations with these host factors. FleA co-localized transiently with ITGB1 at early time points (15-30 min) whereas co-localization with MRC2 was evident from 15 min and persisted up to 4 h.

Although we did not detect proximity between ITGB1 and MRC2 by PLA in FleA-stimulated cells. FleA showed sustained association with MRC2 and later co-localized with LAMP1, consistent with progression toward intracellular compartments. In line with this, PLA signals indicated that MRC2-LAMP1 proximity increased upon FleA stimulation at later time points, suggesting that FleA may reach LAMP1-positive compartments via an MRC2-associated route, rather than through direct interaction with LAMP1.

We propose a model in which ITGB1 acts as an early cellular partner of FleA, promoting subsequent recruitment of MRC2 to the plasma membrane, where it engages the lectin more stably. MRC2 could then mediate intracellular routing of FleA toward vesicular compartments to acquire the lysosomal marker LAMP-1. Entrapment of FleA could be part of the cellular antifungal activity of BECs, preventing conidia from transitioning to a filamentous state.

ITGB1 has been described as a receptor the *A. fumigatus* adhesin CalA (a thaumatin-like protein) (27). Liu *et al*. showed that CalA contributes to epithelial invasion but, similar to FleA in our model, is dispensable for adherence. They also suggested the existence of an additional host cell receptor, since the Δ*calA* mutant displayed reduced invasion of an integrin β1-deficient embryonic stem cell-derived line, compare with the wild-type strain (27).

MRC2 (also known as Endo180/uPARAP) is a member of the macrophage mannose receptor family and is a key collagen receptor involved in collagen internalization and degradation in mesenchymal cells and subsets of macrophages (28). Consistent with a trafficking role, Kjøller *et al*., demonstrating using fluorescently labelled collagen, that MRC2 directed delivery of collagen to vesicular compartments containing LAMP-1 (29). LAMP1 itself has been implicated in conidial intracellular fate in epithelial cells: Wasylnka *et al*., reported that a substantial fraction of internalized conidia co-localized with LAMP1 and that some internalized conidia persist within alveolar epithelial cells (30).

Overall, our findings support an epithelial antifungal response that involves at least two complementary pathways: (i) a PI3K/laminin-332 axis that enhances conidial adhesion, and (ii) a FleA-dependent pathway involving ITGB1 and MRC2, consistent with a role in lectin engagement and intracellular trafficking toward LAMP1-positive compartments.

While our previous work did not reveal a contribution conidial internalization to the antifungal activity of BECs (16), the present data suggest that FleA engagement by ITBG1/MRC2 promotes a route leading to conidia intracellular sequestration and altered fungal development, culminating in reduced filament formation and trafficking toward late endosomes/lysosomes The apparent discrepancy with our earlier conclusions may stem from that we previously assessed internalization primarily through PI3K inhibition, whereas we now show that PI3K chiefly regulates laminin-332 expression, which impacts the earliest step of the epithelial-fungal encounter, adhesion.

Finally, FleA lectin contributes both to the inflammatory response (12) and to epithelial antifungal activity (16). Our results further support FleA as an important virulence factor during early colonization, underscoring the value of developing FleA antagonists to refine our understanding of its role in pathogenesis and to evaluate their potential as therapeutic tools.

## Materials and Methods

### Fungus culture

*Aspergillus fumigatus* DAL strain (CBS 144.89), originally isolated from a human patient with invasive aspergillosis, was maintained on 2% malt agar for one week at room temperature (RT). Conidia were harvested from the slant into 0.1% Tween-20 in phosphate-buffered saline (PBS). The resulting suspension was centrifuged for 10 min at 750 x g, and the pellet was resuspended in PBS/Tween. Conidia were counted using a haemocytometer and adjusted to the desired concentration.

### Bronchial epithelial cells, cell culture and stimulation

Human bronchial epithelial cells (BEAS-2B, ATCC) were maintained at 37°C and 5% CO_2_ and serially passaged in F-12 culture medium (Invitrogen) supplemented with 10% foetal calf serum (FCS) (Eurobio), 1% penicillin and streptomycin (Invitrogen), and 10 mM HEPES in 75-cm^2^ culture flask. Cells were seeded (3 x 10^5^ cells) into 24-well plates 2 days before stimulation. BEAS-2B cells were infected for 15 h with 10^3^ conidia in 300 µL per well of F-12 culture medium containing 10% FCS, without antibiotics, corresponding to a MOI of 0.001. When indicated, BEAS-2B were pretreated for 1 h with PI3K inhibitor, LY294002 (30 µM) or vehicle control (DMSO) before the infection. In separate experiments, BEAS-2B cells were pre-incubated for 1 h recombinant human laminin-332 (BioLamina AB) at 0.1, 1, 10 or 20 µg/mL prior infection. Laminin 332 was dialyzed overnight at 4°C in PBS (Pur-A-Lyser Mini 6000, Sigma) to remove NaN_3_.

### Transcriptomic analysis

#### RNA purification and library construction

Total RNA (2-5 µg), extracted using the NucleoSpin RNAII® kit (Macherey-Nagel, Düren, Germany), was used for polyadenylated mRNAs purification and library construction using the TruSeq RNA Sample Prep Kit v2 (Illumina, #RS-122-2001 and #RS-122-2002, San Diego, CA) according to the manufacturer’s instructions. The nondirectional libraries thus obtained were CTRL led by Bioanalyzer DNA1000 Chips (Agilent Technologies, #5067–1504, Santa Clara, CA) and quantified by spectrofluorimetry (Quant-iT™ dsDNA High-Sensitivity Assay Kit, #Q33120, Invitrogen, Life Technologies, Carlsbad, CA).

#### Statistical analysis

Counts data were analysed using R version 3.0.2 (31) and the Bioconductor package DESeq2 version 1.2.9 (32). Data were normalized using DESeq2 and the default parameters. To estimate dispersion and test for differential expression, we used the default parameters (including outlier detection and independent filtering). The generalized linear model was set with time, inhibitor as well as the time x inhibitor interaction. At each time point, the inhibitor effect was tested using a specific contrast and raw P values were adjusted for multiple testing according to the Benjamini and Hochberg procedure. Genes with an adjusted P value lower than 0.001 were considered differentially expressed (33).

#### Bioinformatic analysis

Sequencing of the 18 samples was performed on the HiSeq 2000 sequencer (Illumina) in single-end mode in order to have approximately 100 million reads of 50 bases per sample. Reads were cleaned of adapter sequences and low-quality sequences using an in-house program (https://github.com/baj12/clean_ngs). Only sequences at least 25 nt in length were considered for further analysis. TopHat (version 1.4.1.1, default parameters, http://ccb.jhu.edu/software/tophat) was used for alignment on the reference genome (hg19). Genes were counted using HTseq-count (parameters: -m intersection-nonempty, -s yes, -t exon) (34),(35). The transcriptomic data have been deposited to the EMBL’s European Bioinformatics Institute with the dataset identifier E-MTAB-16714.

### Microscopic score and galactomannan measurement

As previously described (16), fungal growth was evaluated after 15 h of infection by (i) microscopic scoring of filament formation and (ii) measurement of galactomannan release. For microscopic assessment, filamentation was scored on an arbitrary scale from 1 (no filament) to 6 (maximum level of filament formation). The reference score (3) correspond to filament formation on epithelial cells under contril condition (no inhibitor or additional stimulation). For each experimental condition, 3 wells were used per set of experiments and 2 independent blind observers scored microscopic observations. Galactomannan, a polysaccharide released during *A. fumigatus* growth, was used as an additional quantitative readout of filament formation. Galactomannan antigen levels were measured using a commercially assay (BIO-RAD, Hercules, USA). Supernatants were processed according to the manufacturer’s protocol for serum samples. Results were expressed as a percentage of the values for the control.

### Cell transfection with siRNA

BEAS-2B cells were seeded at 2×10^5^ cells per well in 24-well plates and transfected with siRNA for 72 h. Specific siRNAs (Ambion) were used targeting ITGB1 (20 nM), LAMB3 and LAMC2 (10 nM), MRC2 (20 nM), and LAMP1 (20 nM). After 72 h of transfection, cells were infected with *A. fumigatus* conidia for 15 h. Culture supernatants were then collected and conidia filamentation was quantified by measuring galactomannan release.

### RNA extraction and RTqPCR

BECs were lysed and total RNA was extracted using the Nucleospin miRNA kit (Macherey Nagel, Germany) according to the manufacturer’s instructions. RNA concentrations were measured with a Nanodrop spectrophotometer (Thermo Scientific, Wilmington, USA) and samples were stored at −20°C until use. Reverse transcriptionwas was performed from 1µg RNA using a 2X RT Master Mix on a ProFlex thermocycler (Applied Biosystems, ThermoFischer), with the following program: 25°C for 10 min; 37°C for 2 h and 85°C for 5 min. Quantitative PCR was carried out using 2 µL of cDNA, 5 µL 2× TaqMan mix (Low ROX), 2.5 µL of RNase-free H_2_0 and 0.5 µL of the TaqMan probe: GAPDH (Hs02786624_g1); ITGB1 (Hs01127536_m1); MRC2 (Hs00195862_m1); LAMP1 (Hs00931461_m1); LAMB3 (Hs00165078_m1); LAMC2 (Hs01043717_m1) (Applied Biosystem). Quantitative PCR was done on a Step One Plus PCR System (Applied Biosystem) using 40 cycles with initial denaturation of 95 °C for 2 min, followed by 95 °C for 10 s and 60 °C for 20 s. Relative gene expression was calculated using the ΔCt method with GAPDH as the endogenous control and the Relative Quantity (RQ) was determined by the 2^-ΔΔCT^ method.

### FleA purification and labelling

FleA was expressed and purified as previously described (16). The final buffer exchange was performed either into PBS for labelling and storage at 4°C or into Milli-Q water prior to lyophilization and storage at −80°C. FleA was biotinylated using biotinamidohexanoyl-6-aminohexanoic acid N-hydroxysuccinimide ester (B3295, Sigma-Aldrich/ Merk) in PBS according to manufacturer’s instructions. For fluorescent labelling, FleA was conjugated to FITC (F3651, Sigma-Aldrich/ Merk) following the supplier’s recommendations and stored protected from light in PBS at 1 mg/ml. For long-term storage, biotinylated and FITC-labelled FleA were kept at −20°C.

### Conidial adhesion

To evaluate the respective role of FleA and laminin-332 in conidial adhesion to cells, BECs were pre-incubated for 1 h with either exogenous laminin-332 (20 µg/mL) or FleA (2 µM) and then infected with conidia for 2 or 4 h. Supernatants were collected, plated onto Malt Petri dishes, and incubated overnight at 37°C. The colony-forming unit (CFU) were counted to quantify non-adherent conidia. The number of adherent conidia was then calculated by subtracting non-adherent CFU from the initial inoculum.

### Immunofluorescence and confocal microscopy

BEAS-2B cells were cultivated on glass slides (12mm; 1.5H) in 24 well plates, with a concentration of 5.10^5^ cells/well/mL and incubated for 24 h at 37°C. Cells were incubated with FITC-labeled FleA (0.1µM) or left untreated for the indicated at 37°C. After stimulation, cells were fixed with 4% paraformaldehyde (PFA) for 15 min at RT, washed three times with PBS, and blocked in PBS/BSA 1% for 1 h at RT. Cells were incubated overnight at 4°C with the following primary antibodies diluted in PBS/BSA 1% : anti-ITGB1 (rabbit IgG, Proteintech, USA; 1:1000), anti-MRC2 (mouse IgG, R&D Systems, USA; 1:60) and/or anti-LAMP1 (rabbit IgG; 1:400, or mouse IgG1, 1:100; Cell Signaling, USA). The next day coverslips were washed three times with PBS, and incubated for 1 h at RT with secondary antibody (1:2000) : anti-rabbit IgG Alexa Fluor 594 and anti-mouse IgG AlexaFluor 647 (Invitrogen, USA). Nuclei were counterstained with DAPI. Coverslips were mounted using ProLong Glass Antifade Mountant (Invitrogen, USA) and stored at 4°C. Confocal microscopy was performed using Olympus Fluoview3000.

#### Proximity ligation assay for detection of protein-protein interaction

BEAS-2B cells were seeded onto 12-mm glass coverslips (1.5H) in 24 well plates, at 5×10^5^ cells/well/mL and incubated for 24 h at 37°C. Cells were incubated with FITC-labelled FleA (0.1µM) or left unstimulated for the indicated times at 37°C. Cells were then fixed with 4% of PFA for 15 min at RT, washed three times with PBS, and blocked for 1 h at 37°C in humidified chamber using Duolink Blocking Solution (Duolink® Starter Kit, Sigma-Aldrich, USA). Primary antibodies: were diluted in Duolink® Antibody Diluent and applied overnight at 4 °C: anti-ITGB1 (rabbit IgG, Proteintech; 1:1000), anti-MRC2 (mouse IgG, R&D Systems, USA; 1:60) and/or anti-LAMP1 (rabbit IgG, 1:400 or mouse IgG1, 1:100, Cell Signaling, USA)). Coverslips were washed twice for 5 min with Duolink® Wash Buffer A. PLA probes (anti-rabbit PLUS and anti-mouse MINUS, or the appropriate combination) were diluted in Duolink® Probe Diluent, applied to coverslips, and incubated for 1 h at 37 °C in a humidified chamber. After washes in Wash Buffer A, ligation was performed using the Duolink® Ligation Solution for 30 min at 37 °C, followed by rolling-circle amplification with the Duolink® Amplification Solution for 100 min at 37 °C. Coverslips were then washed twice for 10 min in Duolink® Wash Buffer B and briefly rinsed for 1 min in 0.01× Wash Buffer B. Finally, coverslips were mounted using Duolink® Mounting Medium containing DAPI and stored at −20 °C.

#### Image processing and quantitative analysis

Image processing and quantitative analysis were performed using Fiji/ImageJ (v1.54p or later) with a custom macro designed to automate nuclear counting, puncta detection and fluorescence measurements (36). For each field, a maximum-intensity projection was generated along the z-axis. Nuclei were counted using the *StarDist 2D* model “Versatile (fluorescent nuclei)” (37). Fluorescence quantification was then performed on the red and far-red channels, after thresholding with using user-defined lower limits (*R* and *RL*) and analyzed with the *Analyze Particles* function to measure signal area and intensity. Fluorescent puncta (“dots”) were detected in the far-red channel using the *Find Maxima* function, with the *prominence* parameter (*DOT*) controlling detection sensitivity. All processed projections, masks, and quantitative results were automatically analyzed (**Supplementary file 1**).

### Co-precipitation of cellular receptor of FleA

The co-precipitation was realized by using a commercially available kit (Pierce-MS compatible Magnetic IP Kit, Thermofisher). To obtain 1 to 5 mg of proteins per experimental condition, one 75-cm^2^ culture flask was used per experimental condition, realised in triplicate. BEAS-2B cells were pre-incubated with fungal lectin FleA or biotinylated fungal lectin FleA (2 µM) in 5 mL of PBS during 1 h at 4°C. Cells were lysed with IP-MS Cell Lysis Buffer containing proteases inhibitors for 10 min, on ice. Cell debris were removed by centrifugation at 13,000 x *g* for 10 min. The supernatant was incubated with streptavidin magnetic beads, during 1 h at RT under agitation. Beads were collected with a magnetic stand. After washes, biotinylated lectin and the co-precipitated proteins were eluted with IP-MS Elution Buffer. To allow the proteomic analysis by LC-MS/MS, the eluate was dried in a speed vacuum concentrator.

#### SDS-PAGE

Lyophilized eluates were resuspended in 20 µl Laemmli buffer (62.5 mM Tris/HCl pH 6.8; 2% SDS; 10% glycerol; 0.01% Bromophenol blue; 25 mM DTT), sonicated for 5 min in an ultrasonic bath and then heated at at 60 °C for 5 min with gentle agitation. Samples were centrifuged at 14,000 rpm for 2 min to pellet the residual beads and supernatants were loaded onto a 10 % SDS-PAGE gel. Electrophoresis was performed for a short run (∼1 cm). Gels were fixed overnight in 30% ethanol / 5% acetic acid with moderate shaking, then washed twice with 20% ethanol and 10% ethanol (10 min each), followed by two washes in Milli-Q water (5 min each). For silver staining, gels were sensitized for 1 min in 0.02% sodium thiosulfate, briefly rinsed twice with Milli-Q water and incubated in 0.2 % silver nitrate for 30 min with a with gentle shaking, protected from the light. Gels were then rinsed twice (15 sec each) in Milli-Q water and developed in 3% sodium carbonate containing 0.00125% sodium thiosulfate and 0.03% formalin until bands reached the desired intensity. Development was stopped by replacing the developer with 5% acetic acid for 15 min. Gels were washed and stored in Milli-Q water. For downstream proteomics, entire lanes were excised and cut into ∼1 mm³ pieces for in-gel protein tryptic digestion.

#### In-gel digestion

Excised gel bands were destained twice (2 x 5 min) in a destaining solution (15 mM potassium ferricyanide and 50 mM sodium thiosulfate) and then washed (4 x 5 min) in ultra-pure water. Two additional washes (2 x 10 min) were performed with a 70 % acetonitrile (CAN) solution. Gel pieces were incubated 10 min in pure ACN and air-dried for 30 min. Proteins were reduced and alkylated by incubation of gel pieces in 10 mM DTT, 50 mM ammonium bicarbonate for 30 min at 56 °C followed by incubation in 50 mM iodoacetamide, 50 mM ammonium bicarbonate for 30 min at RT in darkness. Gel pieces were washed twice (2 x 10 min) with 50 mM ammonium bicarbonate, incubated 10 min in pure ACN and air-dried for 30 min. Gel pieces were rehydrated on ice in 50 mM ammonium bicarbonate/5% ACN containing trypsin (3.5 ng/μl; Promega) and digested overnight at 37°C. Supernatants were collected and gel pieces were rinsed twice (2 x 10 min) with 0.1% trifluoroacetic acid and 60% ACN in an ultrasonic bath to extract residual peptides. Peptides were dried out in a vacuum centrifuge, and resuspended in 0.1% formic acid/2% ACN.

#### StageTips

Automated peptide desalting was performed with the DigestPro MSi station (CEM) using home-made StageTips (3 layers of 3M Empore C18 Disc stacked in a 10 µl tip). StageTips were rehydrated with 100% methanol and equilibrated with 0.5 % acetic acid. Peptides were diluted 3.5-fold in 0.5 % acetic acid solution before being loaded on StageTips. StageTips were then washed with 0.5 % acetic acid and peptides were eluted with 80 % CAN/0.5% acetic acid. Eluted peptides were dried out in a vacuum centrifuge and resuspended in 0.1% formic acid/2% ACN.

#### LC-MS/MS

Peptides were analyzed with a nanoElute UHPLC (Bruker) coupled to a timsTOF Pro mass spectrometer (Bruker). Peptides were separated on an Odyssey RP-C18 analytical column (25 cm, 75 μm i.d., 120 Å, 1.6 µm IonOpticks) at a flow rate of 400 nL/min, at 50°C, with mobile phase A (ACN 2 % / FA 0.1 %) and B (ACN 99.9% / FA 0.1%). A 30 min elution gradient was run from 0% to 3% B in 1 min, 3% to 15 % B in 17 min then 15% to 23% B in 7 min and 23% to 32% B in 5 min. MS acquisition was run in DDA mode with PASEF. Accumulation time was set to 100 msec in the TIMS tunnel. Capillary voltage was set to 1,6 kV, mass range from 100 to 1700 m/z in MS and MS/MS. Dynamic exclusion was activated for ions within 0.015 m/z and 0.015 V.s/cm² and released after 0.4 min. Exclusion was reconsidered if precursor ion intensity was 4 times superior. Low abundance precursors below the target value of 20,000 a.u. and intensity of 2,500 a.u. were selected several times for PASEF-MS/MS until the target value was reached. Parent ion selection was achieved by using a two-dimensional m/z and 1/k0 selection area filter allowing the exclusion of singly charged ions. Total cycle time was 1.29 sec with 10 PASEF cycles.

#### Semi-quantitative analysis (spectral counting)

Raw data were processed using Data Analysis 5.1 (Bruker) to generate mgf files. Mgf files were processed with X!Tandem pipeline 0.2.36 using X!Tandem engine 2017.2.1.4. The search parameters were as following: mass tolerance of 25 ppm for MS1 and MS2, carbamidomethyl as fixed modification and oxidation (on M residue) as variable modifications; 1 missed cleavage allowed. Searches (normal and decoy) were performed against the combination of 2 databases: UniProt *Homo sapiens* database (2019-04 release) including the supplementary sequence of Fucose-specific lectin FleA from *A. fumigatus* (UniProt ID = A0A0J5Q677_ASPFM) and a contaminants database. The identification p-values of peptides and proteins were adjusted to get corresponding FDR < 1%, with a minimum of 2 peptides per protein. Proteins from contaminants database were removed. The specific spectral counts of significantly identified proteins were used to compare protein abundances between the different conditions. To highlight candidate proteins, protein list was filtered to retain only proteins present at least in 2 from 3 biotinylated lectin samples and not detected in all control samples. The mass spectrometry proteomics data have been deposited to the ProteomeXchange Consortium via the PRIDE partner repository with the dataset identifier PXD074553 (38).

### Surface plasmon resonance (SPR)

All SPR experiments were performed on a Biacore X100 instrument (Cytiva) in HBS running buffer (10 mM Hepes/NaOH, pH 7.5, 150 mM NaCl, 0.05% Tween 20, 3 mM EDTA) at a flow rate of 20 μl min^-1^ at 25°C. Streptavidin (100 µg/mL) was first immobilized on research grade CM5 chip on the first two flow channels to 3500 resonance unit (RU) and then biotinylated FleA was immobilized on flow channel two to 840 RU following the amine coupling procedure. Single cycle kinetics were performed where ITGB1 was diluted in running buffer prior injection using the same concentration range as in BLI (1.85, 5.5, 16.6 and 50 nM) with a contact time of 270 s and a dissociation time of 80 seconds followed by a regeneration step of 30 s at 30 µl/min with 50 mM NaOH. Binding was measured as resonance units over time after blank subtraction. Data were evaluated by using the Biacore X100 evaluation software, version 2.0. Dissociation constants were determined by plotting response at equilibrium (Req, 10 s before the end of injection) against analyte concentration and the apparent K_D_ was obtained from the steady state analysis using 1:1 association fit.

### Bio-layer interferometry (BLI)

The BLI experiments were performed using an Octet RED96 at 25°C using a 200 µl well volume and in PBS pH 7.4 (Na_2_HPO_4_ 10 mM, KH_2_PO_4_ 1.76 mM, KCl 2.7 mM, NaCl 137 mM) as buffer. The data were collected with ForteBio DataAcquisition9, analyzed and fitted with ForteBio DataAnalysis9 after subtraction of the reference sensor data and adjustment of the baseline to zero. Kinetic titration series were performed and apparent K_D_ was determined from steady state analysis using ForteBio Data Analysis 9.0 and a 1:1 association fit.

#### Streptavidin sensors

Biotinylated FleA was prepared at a concentration of 100 ng/µL for immobilization on streptavidin (SA) sensors (Sartorius). Sensors were incubated in PBS for 10 min before the loading procedure which consisted of: baseline-PBS: 60 s, association 800 s, dissociation 120 s prior equilibration 120 s and loading 600 s. Since we could not fully regenerate the sensors, single cycle kinetics protocol was used to measure the binding of ITGB1 (Sino Biological reference 10587-H08H1) to FleA using a 1000 to 15.63 nM concentration range prepared from two-fold serial dilution of ITGB1. Duplicate was performed with baseline 60 s), association 800 s, dissociation 120 s. Reference sensors were not loaded with biotinylated FleA and were dipped into the same solutions as sensors with immobilized FleA.

#### Ni-NTA biosensors

Another experiment was performed by immobilising ITGB1 or MRC2 (Antibodies online, reference ABIN7490306, residues 31-530) thanks to their N-terminal Histag on NTA sensors (Sartorius). Immobilization: 200 *μ*L of the His-tagged ITGB1 or MRC2 was prepared at 20 µg/mL in PBS pH 7.4 and added to the 96 well plate. Sensors were regenerated with Ni2+ according to manufacturer’s protocol before loading. The kinetic serie sequence used was as follows: (1) baseline (PBS, 80 s), (2) loading (His-tagged protein, 500 s), (3) baseline (PBS, 200 s), (4) association (FleA, 600 s), (5) dissociation (PBS, 600 s) and (6) regeneration (fucose 1m, 40 s) with steps 4-6 repeated for each concentration without regeneration (3.9, 16.65, 65.5, 250, 1000, 1500 nM). Reference sensors and analysis on duplicate experiments were performed as described above.

### Statistics

Data are presented as the mean value ± standard error of mean (SEM). Statistic tests were performed using Prism 7.0 software (GraphPad Software). Differences between two groups were tested using unpaired t-test. ANOVA was performed to compare quantitative variables across groups. To correct for multiple testing, we used Bonferroni’s method, p < 0.05 was considered statistically significant.

## Acknowledgements

This work was supported by a grant from French cystic fibrosis non-profit organizations, Vaincre la Mucoviscidose and Association Gregory Lemarchal (RF20160501616, RF20170501940, RF20180502240, RF20190502450).

The ICMG (UAR 2607) is acknowledged for providing access and support to the Octet RED96 Instrument on the PCI platform for BLI experiments. We would like to thanks Luca Pisapia and Marguerita Duca for performing the BLI measurements and Océane Ricloux for help in BLI data analysis. Biomics Platform, C2RT, Institut Pasteur, Paris, France, is supported by France Génomique (ANR-10-INBS-09) and IBISA.

## Author contributions

**Conceptualization:** N. Millet, V. Balloy, J. Bigot

**Data curation:** N. Millet, V. Balloy, J. Bigot

**Formal analysis:** N. Millet, A. Varrot, C. Pionneau, H. Varet, R. Morichon, V. Balloy, J. Bigot

**Funding acquisition**: Viviane Balloy

**Investigation**: N. Millet, A. Moreau, M. Tarizzo, L. Marti, A. Varrot, E. Gillon, N. Richard, C. Pionneau, S. Chardonnet, H. Varet, R. Morichon

**Methodology:** N. Millet, A. Varrot, C. Pionneau, S. Chardonnet, H. Varet, R. Morichon, V. Balloy, J. Bigot

**Project administrations**: N. Millet, V. Balloy, J. Bigot

**Resources:** N. Millet, V. Balloy, J. Bigot

**Software:** A. Varrot, C. Pionneau, H. Varet, R. Morichon

**Validation:** N. Millet, A. Varrot, C. Pionneau, H. Varet, R. Morichon, V. Balloy, J. Bigot

**Visualization:** N. Millet, V. Balloy, J. Bigot

**Writing - original draft:** N. Millet, V. Balloy, J. Bigot

**Writing – review and editing:** N. Millet, A. Moreau, M. Tarizzo, L. Marti, A. Varrot, E. Gillon, N. Richard, C. Pionneau, S. Chardonnet, H. Varet, R. Morichon, J. Guitard, L. Guillot, V. Balloy, J. Bigot

**Supplementary figure 1: Measurement of galactomannan released by *A. fumigatus* after infection of BECs incubated with lectin FleA biotinylated or not.**

Measure of galactomannan released in supernatant of BECs pre-incubated with lectin FleA or biotinylated lectin FleA (2 µM) for 1 h before infection with *A. fumigatus*. Data are expressed as percentage of control (*A.fumigatus* without epithelial cells) and presented as mean ± SD, n = 4 independent experiments. *p < 0.05, <** p < 0.01, **** p < 0.0001 (ANOVA test followed by Bonferroni’s multiple comparison test).

**Supplementary figure 2: Effect of siRNA on expression**

**(A)** *ITGB1*mRNA expression in BEAS-2B cells treated for 48 h with siRNA Ctrl or siRNA ITGB1 or siRNA ITGB1+MRC2. (**B**) *MRC2*mRNA expression in BEAS-2B cells treated for 48 h with siRNA Ctrl or siRNA MRC2 or siRNA ITGB1+MRC2. (**D**) *LAMP1*mRNA expression in BEAS-2B cells treated for 48 h with siRNA Ctrl or siRNA LAMP1. Data are expressed as % of control and presented as mean ± SD, n = 4-5 independent experiments. *p < 0.05, ** p < 0.01 (ANOVA test followed by Bonferroni’s multiple comparison test)

**Supplementary Table 1: List of genes significantly modulated in BECs treated with vs without PI3 kinase inhibitor and infected by *A. fumigatus* during 0, 2 and 4 h.**

Genes with a P adjusted value < 0.001 were considered differentially expressed.

**Supplementary Table 2: List of genes significantly down-regulated in BECs treated with vs without PI3 kinase inhibitor and infected by *A. fumigatus* during 0, 2 and 4 h.**

The genes selected with a Log2Foldchange < 0.66 and expression < 50 reads

**Supplementary Table 3: List of proteins coprecipitated with FleA.**

Identification by Tandem mass spectrometry of candidate proteins coprecipitated with biotinylated FleA. The identification p-values were adjusted to get corresponding FDR < 1%, with a minimum of 2 peptides per protein.

